# Environmental pheromone and endocrine signals correct heterochronic developmental phenotypes caused by insufficient expression of *let-7* family microRNAs in *C. elegans*

**DOI:** 10.1101/551606

**Authors:** Orkan Ilbay, Victor Ambros

## Abstract

Adverse environmental conditions can affect rates of animal developmental progression and lead to temporary developmental quiescence (diapause), exemplified by the dauer larva stage of the nematode *Caenorhabditis elegans*. Remarkably, patterns of cell division and temporal cell fate progression in *C. elegans* larvae are not affected by changes in developmental trajectory. However, the underlying physiological and gene regulatory mechanisms that ensure robust developmental patterning despite substantial plasticity in developmental progression are largely unknown. Here, we report that diapause-inducing environmental pheromone and endocrine signals correct heterochronic developmental cell lineage defects caused by insufficient expression of *let-7* family microRNAs in *C. elegans.* Two conserved endocrine signaling pathways, DAF-7/TGF-β and DAF-2/Insulin, that confer on the larva diapause/non-diapause alternative developmental trajectories, interact with the nuclear hormone receptor, DAF-12, to initiate and regulate a rewiring of the genetic circuitry controlling temporal cell fates. This rewiring includes: 1) repression of the DAF-12 ligand-activated expression of *let-7* family microRNAs, and 2) engagement of a novel ligand-independent DAF-12 activity to downregulate the critical *let-7* family target Hunchback-like-1 (HBL-1). This alternative HBL-1 downregulation program is responsible for correcting *let-7* family insufficiency phenotypes and it requires the activities of certain heterochronic genes, *lin-46, lin-4* and *nhl-2,* that are previously associated with an altered genetic program in post-diapause animals. Our results show how environmental pheromones and endocrine signaling pathways can coordinately regulate both developmental progression and cell fate transitions in *C. elegans* larvae under stress, so that the developmental schedule of cell fates remains unaffected by changes in developmental trajectory.

## INTRODUCTION

Despite the vast complexity of animal development, developmental processes are remarkably robust in the face of environment and physiological stresses. Multicellular animals develop from a single cell through a temporal and spatial elaboration of events that include cell division, differentiation, migration, and apoptosis. Early developmental cell lineages rapidly diverge functionally and spatially, and continue to follow distinct paths towards building diverse parts of the animal body. Marvelously, the sequence and synchrony of these increasingly complex programs of cell fate progression are precisely coordinated, regardless of various environmental and physiological stresses that the animal may encounter in its natural environment.

The nematode *C. elegans* develops through four larval stages, each of which consists of an invariant set of characteristic developmental events ^1^. During larval development, stem cells and blast cells divide and progressively produce progeny cells with defined stage-specific fates. The timing of cell fate transitions within individual postembryonic cell lineages is regulated by genes of the heterochronic pathway, whose products include cell fate determinant transcription factors, as well as microRNAs (miRNAs) and other regulators of these transcription factors ^2,3^. In mutants defective in the activity of one or more heterochronic genes, the synchrony between cell fates and developmental stages is lost in certain cell lineages, which results in dissonance in the relative timing of developmental events across the animal and consequently morphological abnormalities.

During *C. elegans* larval development, lateral hypodermal stem cells (‘seam cells’) express stage-specific proliferative or self-renewal behavior (Figure 1A). Particularly, whilst seam cells divide asymmetrically at each larval stage (L1-L4), giving rise to a new seam cell and a differentiating hypodermal (hyp7) cell, at the L2 stage, certain seam cells undergo a single round of symmetric cell division, resulting in an increase in the number of seam cells on each side of the animal from ten to sixteen. This L2-specific proliferative cell fate is driven by a transcription factor, HBL-1, which specifies expression of the L2 cell fate, and also prevents the expression of the L3 cell fates ^4,5^. Therefore, in order to allow progression to L3 cell fates, HBL-1 must be downregulated by the end of the L2 stage. If HBL-1 is not properly downregulated, for example in mutants defective in upstream regulatory genes, seam cells inappropriately execute L2 cell fates at later stages, resulting in an enlarged and developmentally retarded population of seam cells in adult worms. Three *let-7* family miRNAs *(mir-48/84/241)* are redundantly required for proper temporal downregulation of HBL-1 ^6^. Larvae lacking all three *let-7* family miRNAs reiterate L2 cell fates in later stages of development. The degree of reiteration, hence the severity of the phenotype, varies depending on genetic and environmental factors ^7^, and can be quantified by counting the number of seam cells in young adult worms.

**Figure 1.**
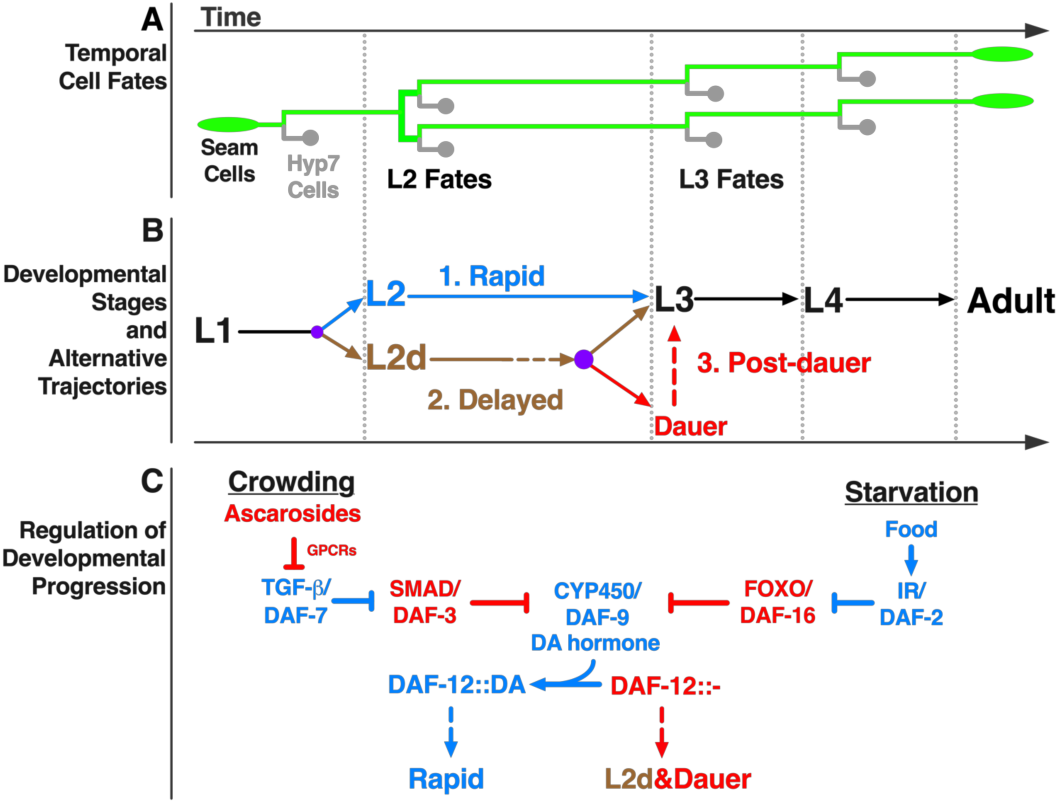
Temporal fates of hypodermal seam cells are robust against changes in developmental trajectory induced by crowding or starvation. **A)** Lineage diagram showing temporal (stage-specific) hypodermal seam cell fates. Seam cells (green) divide asymmetrically at each larval stage, renewing themselves while giving rise to hyp7 cells (gray). At the L2 stage, seam cells undergo a single round of symmetric cell division, resulting in an increase in their number. **B)** Developmental stages and three distinct developmental trajectories: 1. Continuous, unipotent, and rapid progression define the L2 trajectory (blue); 2. Continuous but bipotent and delayed progression define the L2d trajectory (brown), 3. Developmental progression is interrupted by a diapause in the dauer-interrupted or post-dauer trajectory (brown followed by red). Time axis indicates the order of events in time (not proportional to absolute time). Lateral dotted lines indicate the molts between stages. Purple dots represent decision points between different trajectory options. **C)** Regulation of developmental progression. Under favorable conditions dafachronic acid (DA) hormone is abundant and DA-bound DAF-12 promotes rapid development. Crowding or starvation induces dauer formation by repressing TGF-β/DAF-7 signals or insulin signaling (IR/DAF-2: insulin receptor), respectively. Activated effectors of these signaling pathways (DAF-3 or DAF-16) inhibit the biosynthesis of DA; and the unliganded DAF-12 promotes

*C. elegans* is a free-living nematode whose environment is prone to fluctuations between conditions that are favorable and unfavorable for completion of development ^8^. Under favorable conditions (such as abundant food) *C. elegans* larvae develop rapidly and continuously progress through the four larval stages to the adult (Figure 1B, rapid). However, when the conditions are not favorable (for example, in the face of declining resources owing to high population density), the larva at the end of the L2 stage can elect to enter a developmentally arrested diapause, called the dauer larva, which is non-feeding, stress-resistant, and long-lived ^9^. When conditions improve, the dauer larva resumes development to the reproductive adult (Figure 1B, post-dauer). The DAF-7/TGF-β and DAF-2*/*insulin endocrine signaling pathways are the two major signaling pathways that regulate *C. elegans* dauer larva diapause. These two signaling pathways act in parallel to integrate information about population density and nutritional status by co-modulating the biosynthesis of the dafachronic acid (DA) hormone. DA is the ligand of a nuclear hormone receptor, DAF-12, which opposes dauer formation when it is DA-bound and promotes dauer formation when it is unliganded (Figure 1C)^10^.

The order and sequence of temporal cell fates in the various *C. elegans* larval cell lineages are robustly maintained regardless of developmental trajectory: for example, blast cells properly transition from L2 to L3 fates whether the larva develops rapidly and continuously, or instead traverses dauer diapause, which imposes a lengthy (even months-long) interruption of the L2 to L3 transition (Figure 1A&B). Interestingly, cell fate transition defects of many heterochronic mutants are modified (either suppressed or enhanced) when larval development is interrupted by dauer diapause, suggesting that the genetic regulatory pathways regulating temporal cell fate progression are modified depending on whether the animal develops continuously vs undergoes dauer-interrupted development ^11-13^. The mechanisms by which temporal cell fate specification pathways are modified in association with the dauer larva trajectory are poorly understood, especially with regards to how modifications to the regulatory networks controlling temporal cell fate transitions may be coupled to particular steps in the specification and/or execution of the dauer larva diapause trajectory. Of particular interest is the question of whether and how the dauer-promoting signals that are monitored by L1 and L2d larvae might act prior to dauer commitment to directly modify gene regulatory mechanisms controlling temporal cell fate transitions.

To investigate the impact of dauer-inducing environmental and endocrine signals on the regulatory network controlling temporal cell fate transitions, we employed experimental conditions that induce the dauer formation program, but also efficiently prevent dauer commitment. We call these conditions “L2d-inducing” because the presence of both dauer-inducing and commitment-preventing conditions results in worm populations growing continuously (without dauer arrest) but where all animals traverse the lengthened bi-potential L2d stage (Figure 1B, L2d/delayed) ^14,15^.

We found that L2d-inducing pheromones suppress heterochronic defects caused by insufficient expression of *let-7* family microRNAs, suggesting that these pheromones that enable the dauer life history option also activate a program alternative to *let-7* family microRNAs in controlling stage-specific temporal cell fate progression. We found that the two major endocrine signaling pathways that regulate dauer formation in response to pheromones and food signals, the DAF-7/TGF-β and DAF-2/Insulin respectively, also mediate the effect of these same signals on temporal cell fates under L2d-inducing conditions. Moreover, we identified a previously undescribed ligand-independent activity of the nuclear hormone receptor DAF-12 that is responsible for activating the alternative program of cell fate specification in the L2d. This alternative program is responsible for correcting *let-7* family insufficiency phenotypes and it requires the activities of certain heterochronic genes, *lin-46, lin-4* and *nhl-2,* that are previously associated with an altered genetic program in post-diapause animals. This alternative program associated with L2d is coupled to a previously described reduction in the DAF-12-regulated expression of *let-7* family microRNAs ^16^. Hence, the overall L2d response is a “rewiring” program consisting two major operations: 1) repression of *let-7* family microRNA expression, and 2) activation of an alternative program to downregulate the *let-7* family target (Hunchback-like-1) HBL-1.

Our results show that environmental signals and downstream endocrine signaling pathways are capable of coordinately regulating developmental progression and cell fate transitions in *C. elegans*. We propose that this capability confers elasticity to *C. elegans* development, whereby the proper developmental schedule of cell fates remains unaffected by changes or uncertainties in developmental trajectory.

## RESULTS

We developed three approaches to efficiently uncouple L2 from dauer commitment and thereby produce worm populations developing continuously through the bi-potential pre-dauer L2d phase directly to the L3, without dauer arrest. In the first approach, we employed the pheromone cocktail formula described by Butcher *et al.*^17^, which contains three ascaroside molecules (ascr#2, ascr#3, and ascr#5) that synergistically induce L2d and dauer arrest^17^ (Supplemental figure S1A). At sufficiently high doses, the ascaroside cocktail can induce 100% dauer formation. Previous findings showed that the presence of (high-quality) food can antagonize pheromones and prevent dauer formation^14^. However, it was not clear if the food signals also prevented the L2d. We observed that high quality food, or the presence of dafachronic acid (DA) hormone, could efficiently prevent dauer formation while not preventing the L2d, evidenced by dramatically slowed second stage larval development. Therefore, to obtain animal populations traversing the L2d without committing to dauer arrest, we allowed larvae to develop in the presence of a combination of the ascaroside cocktail along with DA hormone^18^ (Supplemental figure S1B-C). Our second approach also uses the ascaroside cocktail, but to eliminate the need for DA hormone, dauer-commitment defective *daf-12(rh61)* mutant worms are employed. In the third approach, to genetically induce L2d, we combined *daf-12(rh61)* with a temperature-sensitive *daf-7* mutant that mimics the L2d-inducing pheromone conditions, or with a temperature-sensitive *daf-2* mutant that mimics the L2d-inducing starvation conditions.

### L2d-inducing ascarosides reduce the reliance on the *let-7* family microRNAs for proper L2-to-L3 cell fate transition

We found that when *mir-48/84/241(0)* mutant larvae developed through L2d -- induced by a combination of the ascaroside cocktail and the DA hormone -- the extra seam cell phenotype of was substantially (albeit partially) suppressed (Figure 2, row 1 vs 3). To compare the strength of this L2d suppression with the previously described post-dauer suppression of the *let-7* family phenotypes^13^, we used the ascaroside cocktail but this time without the DA hormone. Under these conditions *mir-48/84/241(0)* larvae arrested as dauers; and as described previously ^13^ this resulted in complete suppression of the extra seam cell phenotype in post-dauer adults (Figure 2, row 1 vs 2),. Therefore, the L2d suppression is weaker than the post-dauer suppression (Figure 2, row 2 vs 3), and unlike dauer arrest, L2d-inducing ascarosides do not completely eliminate the need for *let-7* family microRNAs for proper L2-to-L3 cell fate transition. Nonetheless, the partial suppression of the extra seam cell phenotype of *let-7* family microRNAs suggests that the L2d-inducing ascarosides rewire the genetic regulatory pathway controlling temporal cell fate progression in a way to reduce the reliance on the *let-7* family microRNAs for proper L2-to-L3 cell fate transition.

**Figure 2.**
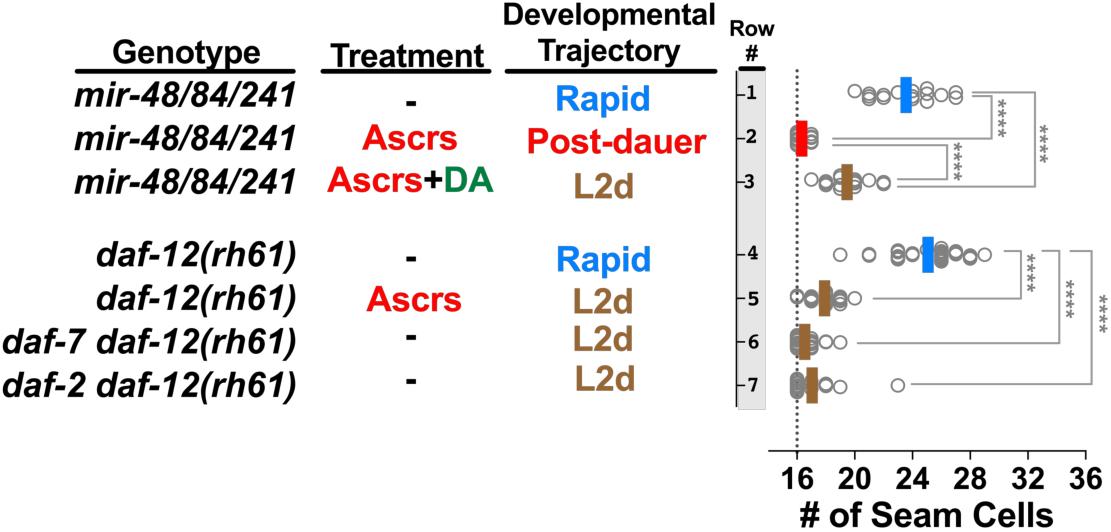
Sensitized genetic backgrounds reveal that L2d-inducing environmental and endocrine signals impact the regulation of temporal cell fates. Number of seam cells in young adult animals cultured under different treatment conditions. Each dot in the plot shows the number of seam cells on one side (left side or right side, observed interchangeably) of a single young adult animal, and solid lines (color code matching the developmental trajectory) indicate the average seam cell number of the animals scored for each condition. Wild-type animals have sixteen seam cells per side, which is indicated by the lateral dotted line. Genotypes indicated in the left column; treatments and corresponding developmental trajectories are indicated in the middle and right columns, respectively. The student’s t-test was used to calculate statistical significance (p): n.s.(not significant) p>0.05, *p<0.05, **p<0.01, ***p<0.001, ****p<0.0001

Each of the individual ascarosides in the cocktail (Ascrs#2,3,5) has been shown previously to be alone sufficient to induce dauer formation, although with reduced potency compared to the combined cocktail ^17,19^. Consistent with their individual capacities to induce L2d and dauer formation, we observed that each ascaroside ascr#2, ascr#3, and ascr#5 alone could suppress the extra seam cell phenotype of *daf-12(rh61)* mutants. In the case of ascr#2 or ascr#3 alone, the suppression was partial, while for ascr#5 alone, the suppression was similar to the full cocktail (Supplemental figure S3A, rows 6 to 10). ascr#5 was the most potent of the three ascarosides in terms of both percent dauer formation of wild type larvae (Supplemental figure S3B) and suppression of the extra seam cell phenotype of *daf-12(rh61)* (Supplemental figure S3A, row 10 vs 8 and 9).

### L2d-inducing ascarosides or L2d-inducing mutations of *daf-7* and *daf-2* suppress heterochronic phenotypes caused by insufficient expression of *let-7* family microRNAs in *daf-12(rh61)* mutants

The *daf-12(rh61)* mutation combines three important properties which makes this mutation uniquely useful for studying the effects of L2d-inducing conditions on the regulation of temporal cell fates. These properties are: 1) *daf-12(rh61)* animals reiterate expression of L2 cell fates owing to reduced (insufficient) *let-7* family levels, 2) *daf-12(rh61)* larvae are unable to execute dauer larvae commitment or arrest ^20^, enabling the use of dauer-promoting conditions to obtain populations of *daf-12(rh61)* animals undergoing an L2d-direct-to-L3 continuous development trajectory; 3) *daf-12(rh61)* animals are insensitive to the DA hormone (due to lack of the DAF-12 ligand binding domain) ^18,20^, and so the levels of *let-7* family microRNAs are expected to be unresponsive to experimentally administered ascaroside s, which are understood to regulate DAF-12 activity by affecting the level of DA ^10^.

We observed that the presence of exogenous ascaroside cocktail during larval development almost completely suppressed the extra seam cell phenotype of *daf-12(rh61)* mutants (Figure 2, row 4 vs 5). To test the possibility that an unexpected elevation in the *let-7* family levels could be responsible for the suppression of the heterochronic phenotypes of *daf-12(rh61)* animals in the presence of the ascaroside cocktail, we quantified the levels of *let-7* family microRNAs in the absence and presence of the ascarosides. No elevation in the levels of these microRNAs in response to the ascaroside cocktail was evident (Supplemental figure S2). Therefore, the suppression of the heterochronic phenotypes of *daf-12(rh61)* mutants in the presence of the ascaroside cocktail is not likely to result from restoration of normal levels of *mir-48/84/241* or an elevation of the other members of the *let-7* family microRNAs.

Similar to the ascaroside cocktail, conditional dauer-constitutive mutants of *daf-7* (mimicking high ascarosides) or *daf-2* (mimicking starvation), almost completely suppressed the extra seam cell phenotype of *daf-12(rh61)* mutants (Figure 2, row 4 vs 6 or 7). These results indicate that genetically induced L2d, whether by activation of the ascaroside response pathway (*daf-7(lf)*), or by activating the starvation response pathway (*daf-2(lf)*), results in an L2d-associated rewiring of the regulatory networks controlling temporal cell fate progression.

### Ascarosides suppress the heterochronic phenotypes of *daf-12(rh61)* via *srg-36/37*-encoded GPCR signaling upstream of DAF-7/TGF-β-DAF-3 signaling

It has been shown previously that induction of dauer formation by ascarosides involves sensing of environmental ascaroside levels by specific G-protein coupled receptors (GPCRs) expressed in chemosensory neurons, wherein they repress DAF-7/TGF-β signals ^21-24^. To test whether these GPCRs were also required for the suppression of the heterochronic phenotypes of *daf-12(rh61)* mutants, we employed mutations of *srg-36* and *srg-37*, which encode GPCRs that are expressed in the ASI neurons and that are redundantly required for perceiving ascr#5 signal in the context of dauer induction^23^. We observed that for *srg-36(0) srg-37(0); daf-12(rh61)* compound mutants, ascaroside (in this case ascr#5) failed to suppress the extra seam cell phenotype *daf-12(rh61)* (Figure 3A, compare row 1 vs 2 with row 3 vs 4). Moreover, we found that the TGF-β signaling effector *daf-3*, which functions downstream of SRG-36/37, is required for the suppression of *daf-12(rh61)* by asaroside (Figure 3B). These results indicate that the same GPCRs that mediate dauer formation in response to asaroside are also required for mediating the effects of asaroside on temporal cell fates, and supports a common pathway for suppression of *daf-12(rh61)* by ascaroside and dauer induction, involving activation of SRG-36/37 GPCRs and the downstream TGF-β effector DAF-3.

**Figure 3.**
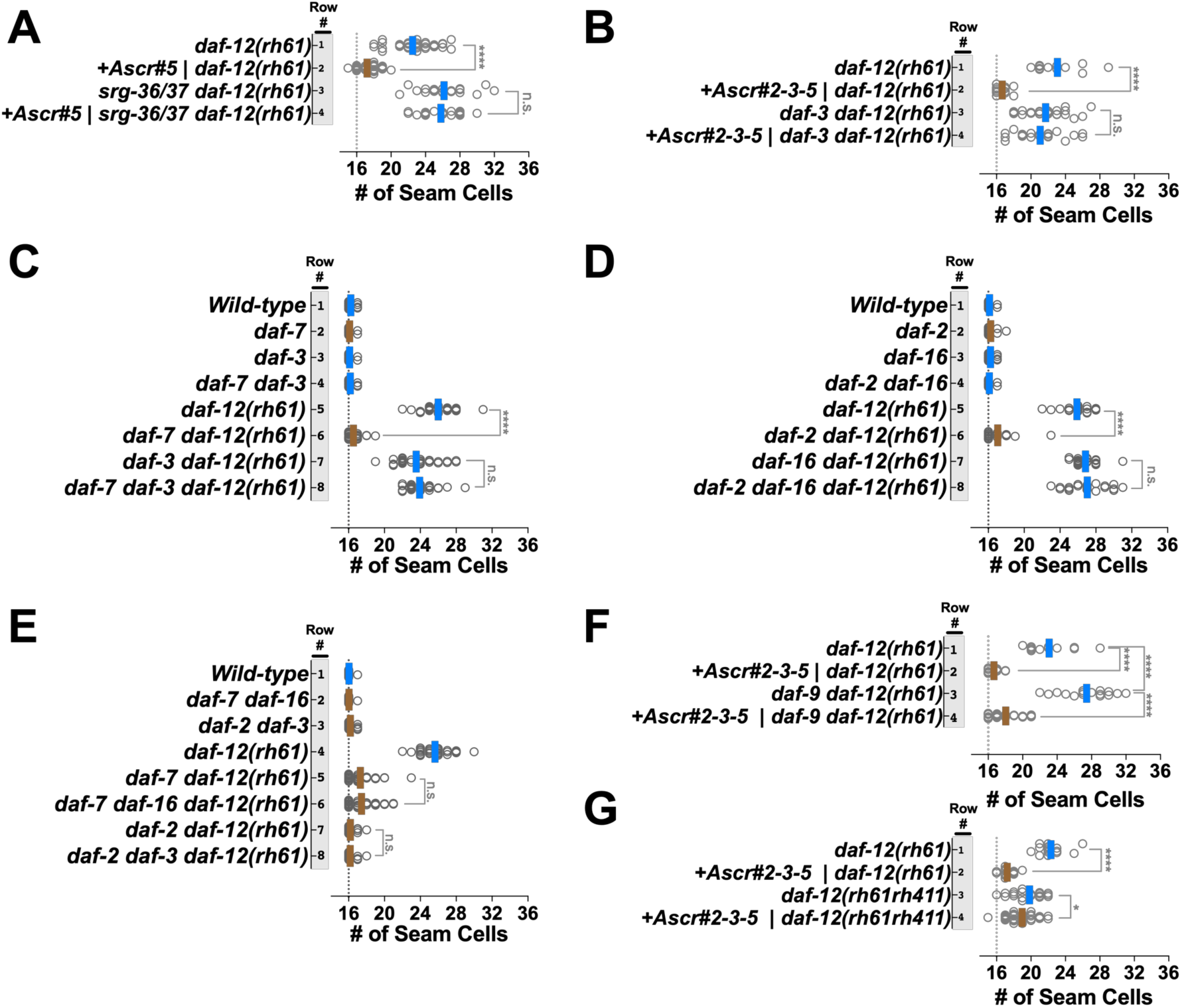
DAF-7/TGF-β and DAF-2/Insulin signaling pathways act in parallel to modulate a ligand-independent activity of DAF-12 that is responsible for correcting heterochronic phenotypes caused by insufficient expression of *let-7* family microRNAs. Number of seam cells in young adult animals of various mutants cultured on ascaroside or control plates (A, B and F. G), or on standard NGM plates (C-E). Each dot in the plots shows the number of seam cells of a single young adult animal, and solid lines indicate the average seam cell number of the animals scored for each condition (blue lines: rapid trajectory; brown lines: L2d trajectory). **A)** Ascarosides suppress *daf-12(rh61)* via *srg-36/37*-encoded GPCR signaling upstream of DAF-7/TGF-β-DAF-3 signaling. **B)** *daf-3* activity is required for suppression of *daf-12(rh61)* by ascarosides. **C-E)** DAF*-*7*/*TGF-β and DAF-2/Insulin signaling pathway act in parallel to mediate the suppression of *daf-12(rh61)*. **F, G)** Ligand-independent activity of *daf-12* is required for the ascaroside-mediated L2d rewiring of the pathways regulating temporal cell fates. The student’s t-test was used to calculate statistical significance (p): n.s.(not significant) p>0.05, *p<0.05, **p<0.01, ***p<0.001, ****p<0.0001.

### DAF-7/TGF-β and DAF-2/Insulin signaling pathway act in parallel to mediate the suppression of the heterochronic phenotypes of *daf-12(rh61)*

As shown above, genetic activation of dauer-inductive signaling by *daf-7(lf)* or *daf-2(lf)* mutations is sufficient for suppression of *daf-12(rh61)* (Figure 2). In the context of dauer formation, DAF*-*7*/*TGF-β primarily mediates ascaroside signaling, and DAF-2/Insulin primarily mediates assessment of nutritional status ^10,21,25^. To determine if the known downstream effectors of DAF*-*7*/*TGF-β and DAF-2/Insulin signaling that mediate dauer formation are also required for mediating the L2d rewiring caused by *daf-7(lf)* or *daf-2(lf)*^26,27^, we generated compound mutants carrying *daf-12(rh61)* in combination with mutations that impair these effectors of the TGF-β or insulin signaling pathways, and determined the number of seam cells in young adults. We found that the downstream effector of the TGF-β signaling pathway, *daf-3*, and the downstream effector of the insulin signaling pathway, *daf-16*, were required for the suppression mediated by the *daf-7(lf)* mutation and the *daf-2(lf)* mutation, respectively (Figure 3C and 3D). These results are consistent with the finding that *daf-3* was also required for the ascaroside-mediated suppression of *daf-12(rh61)* (Figure 3B).

To determine whether the TGF-β and insulin signaling pathways act in parallel to modulate temporal cell fates, we tested for crosstalk between these pathways in context of suppression of *daf-12(rh61)* phenotypes. Specifically, we determined whether *daf-16(lf)* could alter the suppression of *daf-12(rh61)* phenotypes by *daf-7(lf)*, and conversely, whether *daf-3(lf)* could alter the suppression of *daf-12(rh61)* phenotypes by *daf-2(lf).* We found that *daf-16* was not required for *daf-7*-mediated suppression (Figure 3E, rows 5-6), and *daf-3* was not required for *daf-2*-mediated suppression (Figure 3E, rows 7-8), indicating that, similar to their regulation of dauer diapause, the TGF-β and insulin signaling pathways act in parallel in the context of the L2d rewiring of the genetic regulatory pathways controlling larval cell fate progression.

### Ligand-independent activity of *daf-12* is required for the ascaroside-mediated L2d rewiring of the pathways regulating temporal cell fates

Ascaroside (TGF-β) signaling and nutritional status (insulin) signaling converge to induce dauer larva arrest by down regulating DA production and hence reducing the levels of liganded DAF-12. Since we find that dauer-inducing conditions (ascarosides; loss of *daf-7* or *daf-2*) can suppress the heterochronic phenotypes of the DA-insensitive *daf-12(rh61)* mutant, it appears that the TGF-β and insulin signaling pathways may regulate cell fate transitions by repressing a hypothetical DAF-12-independent function of DA. If that were the case, inhibiting or preventing DA production would mimic the effect of ascarosides and suppress the extra seam cell phenotype of *daf-12(rh61).* To test this possibility, we employed genetic ablation of DA production. *daf-9* encodes a CYP450 that is responsible for DA production^18^. Accordingly, *daf-9(lf)* mutants are dauer-constitutive due to lack of DA^18^. To test if ascarosides act by inhibiting DA production during L2d rewiring, we generated double mutants containing *daf-9(lf)* and *daf-12(rh61).* We observed that these double mutants lacking *daf-9* in the *daf-12(rh61)* background had an even stronger extra seam cell fate phenotype than *daf-12(rh61)* mutants (Figure 3F, row 1 vs 3), and that this phenotype was suppressed in the presence of ascarosides (Figure 3F, row 3 vs 4). These results indicate that the ascaroside-induced L2d rewiring does not involve inhibition of DA biosynthesis, nor does rewiring require DA production or *daf-9* activity. The enhancement of the extra seam cell phenotype of *daf-12(rh61)* phenotype in the *daf-12(rh61)*; *daf-9(lf)* double mutant could reflect DAF-12-independent functions of DA or DA-independent functions of DAF-9.

The finding that ascaroside-mediated L2d rewiring did not involve the DA hormone raised the question as to whether the DA receptor, DAF-12 is required for the L2d rewiring. To determine whether *daf-12* is required for ascaroside-induced suppression of retarded seam cell phenotypes, we tested whether ascarosides could suppress the phenotypes of *daf-12(rh61rh411),* a *daf-12* null allele ^20^. *daf-12(rh61rh411)* animals display a milder extra seam cell phenotype than *daf-12(rh61)* ^20^, presumably because of a milder reduction of *let-7* family microRNAs compared to *daf-12(rh61)* ^16^. We observed that the ascaroside conditions that resulted in a very potent suppression of the extra seam cell phenotype of *daf-12(rh61)* animals resulted in only a very modest (albeit statistically significant) suppression of the *daf-12(rh61rh411)* phenotype (Figure 3G, compare changes in the average seam cell in row 1 vs 2 with 3 vs 4). This result suggests that ascaroside-induced L2d rewiring of the pathways regulating temporal cell fates largely requires *daf-12* function, and therefore represents a novel ligand-independent regulation of *daf-12* by TGF-β and insulin signaling.

### Heterochronic genes previously associated with the altered HBL-1 down-regulation program in post-dauer animals are required for the L2d suppression of heterochronic phenotypes caused by insufficient expression of *let-7* family microRNAs

In animals that arrested as dauer larvae and then later resumed development through post dauer larval stages, the genetic programming of temporal cells fates differed substantially from animals that developed continuously ^13^. In particular, proper cell fate progression through dauer larvae arrest and post-dauer development rests on an altered HBL-1 down-regulation program. These differences in HBL-1 down regulation include, 1) reallocation of roles for *lin-4* microRNA and *let-7* family microRNAs, and 2) alterations in the relative impacts of LIN-46 and the microRNA modulatory factor NHL-2 ^13^. For example, animals deficient for *lin-4* exhibited stronger retarded developmental timing phenotypes when traversing developmental arrest followed by post-dauer development compared to animals of the same genotype that developed rapidly and continuously. Similarly, animals carrying loss of function mutations of *nhl-2* or *lin-46* exhibited enhanced retarded phenotypes after post dauer development. *nhl-2* encodes an RNA binding protein that functions as a microRNA positive modulator ^28^, and *lin-46* encodes a protein that acts downstream of the LIN-28 RNA binding protein ^29^. These results suggested that the rewiring of developmental cell fate progression in dauer-traversing larvae involves alterations in the post-transcriptional regulation of HBL-1 expression.

To test whether these previously described genetic requirements for *lin-46, lin-4*, and *nhl-2* in the dauer/post-dauer context^13^, also apply during L2d development, we examined the phenotypes of the relevant mutant strains during development through ascaroside-induced L2d, but in this case without dauer commitment or arrest. We observed that ascarosides failed to suppress the retarded phenotypes of animals that were lacking *lin-46 or lin-4* in combination with *mir-84(lf)* (to provide a sensitized background, blunting the expression of *let-7* family microRNAs), or that were lacking *nhl-2* in the *daf-12(rh61)* background. (Figure 4). Moreover, for doubly-mutant animals carrying both *mir-84(lf)* and *lin-46(lf)* mutations, ascaroside-induced L2d enhanced the retarded phenotypes (Figure 4A), similar to the enhancement reported for *mir-84(lf)*; *lin-46(lf)* animals that developed through dauer arrest and post-dauer development^13^. Similarly, ascarosides failed to suppress the gapped alae phenotype of animals lacking *lin-4* and *mir-84* (Figure 4B), consistent with the previous finding that this phenotype of *lin-4(lf); mir-84(lf)* animals was enhanced for post dauer animals^13^. Lastly, ascarosides failed to suppress the extra seam cell phenotypes of *nhl-2: daf-12(rh61)* animals (Figure 4C), indicating that *nhl-2* is required for ascaroside-mediated suppression of *daf-12(rh61);* analogous to its role in post-dauer enhancement of the retarded phenotypes of *let-7* family microRNAs^13^. These results suggest that the L2d rewiring includes the activation of an alternative HBL-1 downregulation program, which involves *lin-46, lin-4* and *nhl-2*, and that this alternative program accounts for the reduced reliance on *let-7* family microRNAs for proper L2-to-L3 cell fate transition. These results also suggest that the genetic circuitry controlling cell fate progression via HBL-1 in larvae undergoing L2d development is similar to the circuitry associated with dauer larvae arrest, consistent with a rewiring mechanism that is initiated during L2d and augmented during dauer arrest (Figure 4D).

**Figure 4.**
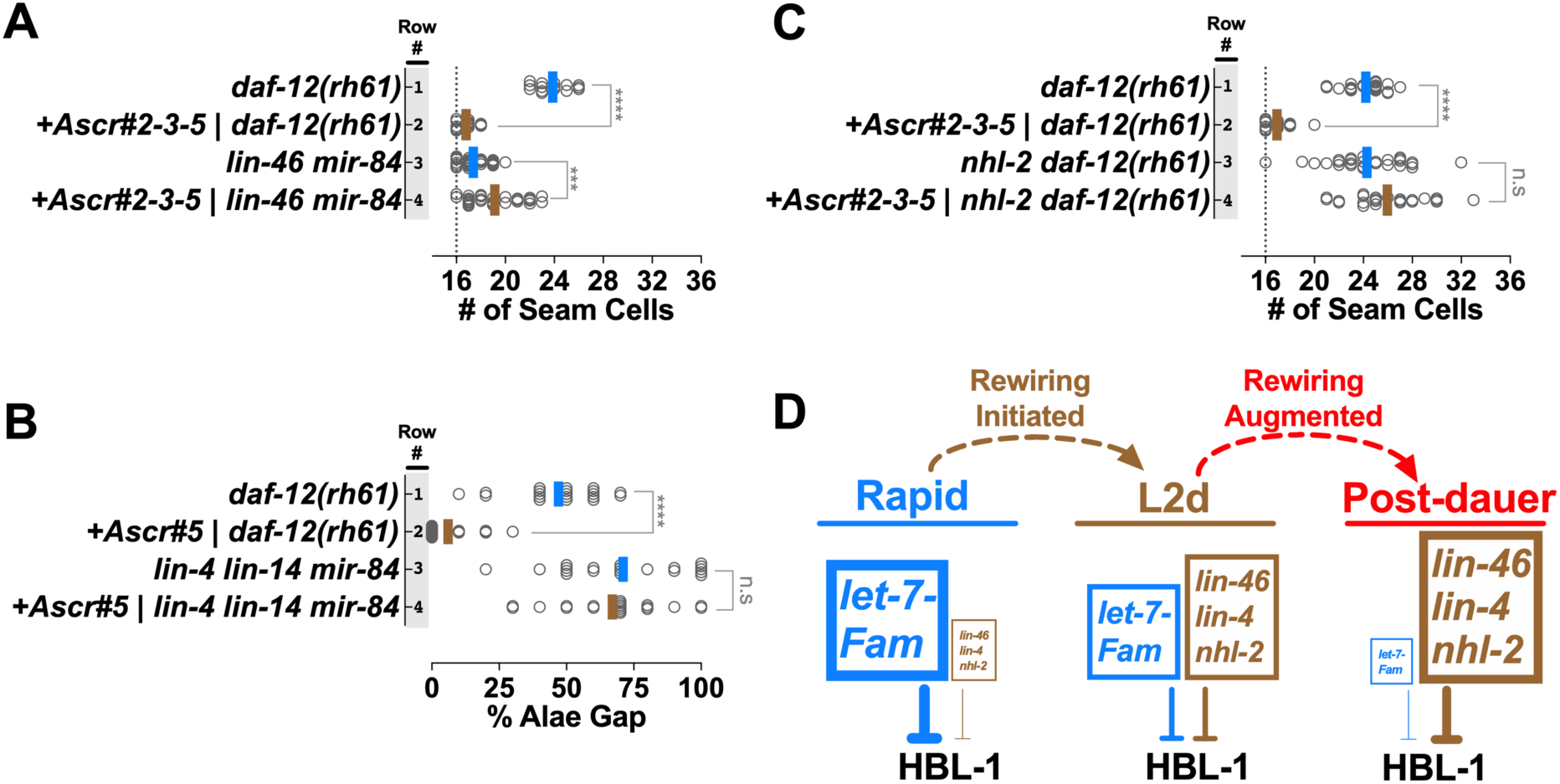
Heterochronic genes associated with the altered HBL-1 down-regulation program in post-dauer animals are required for the L2d suppression of heterochronic phenotypes caused by insufficient expression of *let-7* family microRNAs. **A)** Ascaroside conditions that suppress the extra seam cell phenotype of *daf-12(rh61)* enhance the extra seam cell phenotype of larvae lacking *lin-46* and *mir-84*. **B)** Ascaroside conditions that suppress the gapped alae (a consequence of retarded seam cell development that is manifested in young adults) phenotype do not suppress the gapped alae phenotype of*lin-4; lin-14*; *mir-84* animals. **C)** *nhl-2* activity is required for ascaroside-mediated suppression of *daf-12(rh61)* **D)** A model for the L2d rewiring and its potential augmentation during dauer arrest. Under L2d-inducing conditions, *let-7* family microRNAs are downregulated and also become less important. The reduction in the *let-7* level and importance is coupled to enhanced roles for the heterochronic genes previously associated with the altered HBL-1 downregulation program in post-dauer animals, involving *lin-46, lin-4*, and *nhl-2*. This shift in the reliance on the *let-7* family microRNAs to the reliance on the alternative program for proper HBL-1 down-regulation (hence for proper L2-to-L3 cell fate progression) constitutes the L2d rewiring. In post-dauer animals, consistent with an augmentation of the L2d rewiring program, the reliance on the altered HBL-1 down-regulation program further increases while the *let-7* family microRNAs become dispensable for proper HBL-1 down-regulation. The student’s t-test was used to calculate statistical significance (p): n.s.(not significant) p>0.05, *p<0.05, **p<0.01, ***p<0.001, ****p<0.0001.

## DISCUSSION

Environmental and physiological stress signals can challenge the progression of *C. elegans* larval development, causing the larva to choose one of three distinct alternative developmental trajectories: 1) a rapid and continuous trajectory without the option for dauer arrest, 2) continuous development through an extended, bipotent L2d trajectory, wherein the option for dauer arrest is enabled, but not necessarily selected, and 3) development through the L2d trajectory followed by dauer larva arrest (Figure 1B). Regardless of which trajectory is chosen by the larva, the same sequence of stage-specific cell fates is robustly expressed (Figure 1A). The findings reported here illuminate how genetic regulatory networks that specify larval cell fate progression can accommodate these alternative life histories, and the physiological and environmental stresses that induce them.

Central to the coordination of temporal cell fates and life history choices in *C. elegans* is the nuclear hormone receptor transcription factor DAF-12 ^20^, and its ligand, dafachronic acid (DA)^18^. DAF-12 in the unliganded form is essential for the dauer larva trajectory, while ligand-bound DAF-12 inhibits the dauer larva program. At the same time, DAF-12 and DA control the L2-to-L3 cell fate transitions by regulating the expression of *let-7* family microRNAs, which are required to downregulate HBL-1 and thereby specify the proper timing of expression of L3 cell fates ^6,16,18,30^.

The *let-7* family microRNAs also regulate the abundance of DAF-12 via a feedback loop that has been proposed to help ensure robust coordination of cell fates with dauer arrest^16^. Under favorable conditions, when DA is abundant, DA-bound DAF-12 promotes continuous development, and also activates accumulation of *let-7* family microRNAs during the L2 stage, which in turn attenuate the accumulation of DAF-12, thereby eliminating the dauer option, and also down regulate HBL-1 to enable rapid progression from L2 to L3 cell fates. Conversely, under unfavorable conditions, DA production is low and unliganded DAF-12 promotes the L2d/dauer program and represses the expression of *let-7* family microRNAs. In turn the low level of *let-7* family microRNAs allows the accumulation of DAF-12 during L2d, maintaining the dauer option ^16^.

It is not previously clear how HBL-1 might be down regulated during L2d considering the repressed state of the *let-7* family microRNAs. The results reported here show that during the L2d trajectory, when *let-7* family microRNA levels are low, a previously unknown DA-independent function of DAF-12 is engaged to down regulate HBL-1 and trigger the L2-to-L3 cell fate transition. The key finding pointing to the involvement of DA-independent function of DAF-12 in L2d temporal cell fate control is our observation that the L2d trajectory is conducive to normal cell fate specification for *daf-12(rh61)* (Figure 2), whilst continuous (non-L2d) development results in dramatic retarded heterochronic phenotypes for *daf-12(rh61)* (Figure 2; and ^20^). The L2d suppression of *daf-12(rh61)* phenotypes led us to propose that during wild type L2d, when DA is low, DAF-12 is unliganded and its activity is hence not unlike that of the ligand-binding defective mutant DAF-12(RH61). In this model, we propose that *daf-12(rh61)* animals constitutively run an “L2d-like program” for regulating temporal cell fates, which is discordant with rapid continuous development, but compatible with (and hence phenotypically suppressed by) the environmental and endocrine signals that induce the L2d trajectory. We propose that the L2d program instigated by unliganded DAF-12 includes, in addition to the repression of *let-7* family microRNAs, also the activation of an alternative mechanism of HBL-1 downregulation (Figure 5 and 6), which is responsible for the suppression of *daf-12(rh61)* by ascarosides (Figure 2), and by L2d-inducing mutants of *daf-7* and *daf-2* (Figure 2).

**Figure 5.**
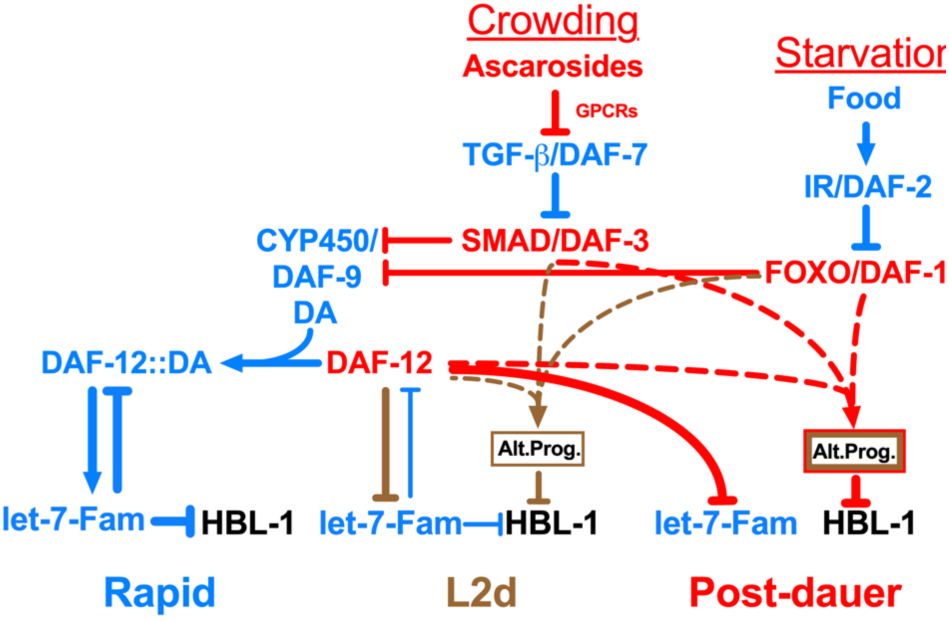
Environmental pheromone and endocrine signals engage DAF-12 to initiate and regulate the rewiring of the HBL-1 down-regulation. In response to crowding and starvation, TGF-β and insulin signaling pathways, respectively, modulate the ligand-dependent DAF-12 activity to repress the transcription of *let-7* family microRNAs, and, at the same time, cooperate with DAF-12 in a ligand-independent manner to activate the alternative HBL-1 downregulation program (Alt. Prog.). The alternative program of L2d and dauer-interrupted trajectories are similar, but the alternative program of dauer-interrupted trajectory is stronger either due to an enhancement of the alternative program of L2d (depicted as thicker brown border line) and/or due to employment of additional factors (depicted as red border line) after the L2d larvae commit to dauer formation.

**Figure 6.**
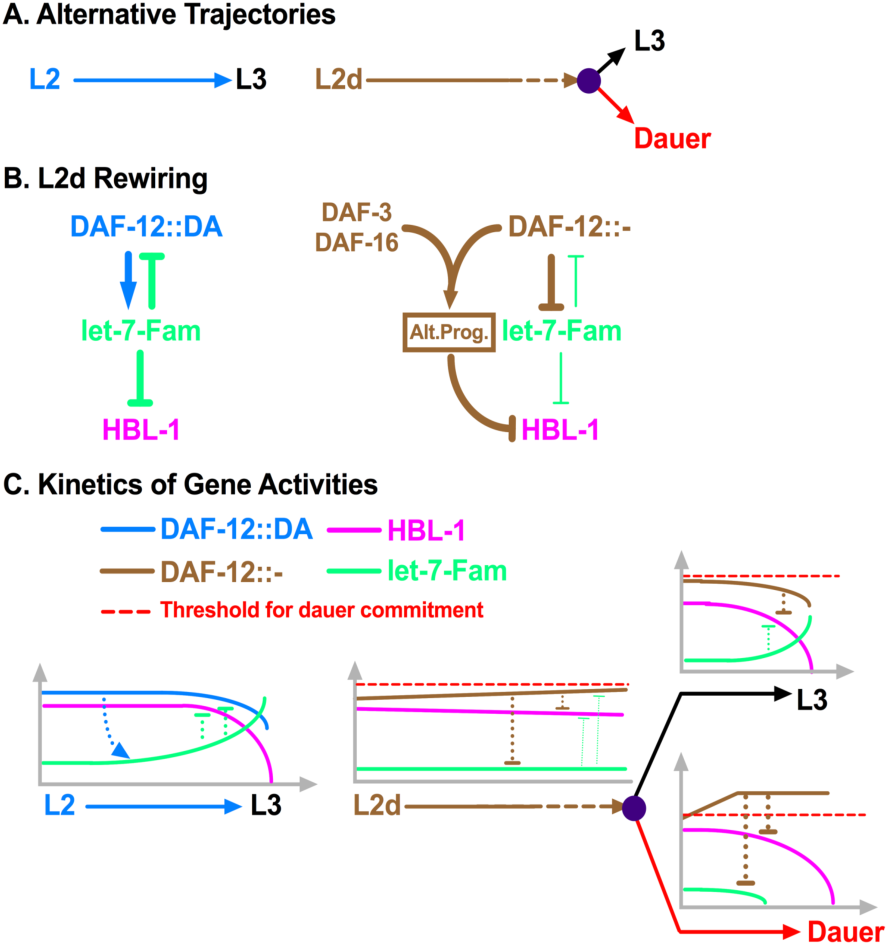
DAF-12 ensures properly delayed but robust HBL-1 down-regulation during L2d-to-L3 transition by coordinating the repression of *let-7* family microRNAs with the activation of the alternative HBL-1 downregulation program. **A)** Alternative trajectories. The L2-to-L3 transition is rapid and deterministic (once committed to the L2 stage, the larvae do not have the dauer option), whereas the L2d-to-L3 transition is slower and bipotential. In both cases, HBL-1 is present throughout the L2/L2d stage but it is downregulated by the beginning of the L3 stage (Supplemental Figure S4). **B-C)** The L2d rewiring and the kinetics of gene activities. During rapid, L2 development, DAF-12 activates the transcription of *let-7* family microRNAs, which in turn, negatively regulate DAF-12, eliminating the dauer option. During slow, bipotential, L2d development, DAF-12 represses *let-7* family microRNAs, which otherwise prevent the accumulation of DAF-12. If the unliganded DAF-12 reaches the threshold, larvae commit to dauer formation, if not, larvae commit to continuous development. While mediating this decision, which necessitates the repression of *let-7* family microRNAs (for maintaining the dauer option) and delaying the downregulation of HBL-1 (for postponing L3 cell fates), DAF-12 cooperates with DAF-3 or DAF-16 to activate the alternative HBL-1 downregulation program (Alt. Prog) to ensure robust HBL-1 downregulation during the L2d-to-L3 transition.

Our findings indicate that the DA-independent function of DAF-12 that modulates temporal cell fates during the L2d can be activated by either of the two upstream transcription factors, DAF-3, which mediates responses to TGF-β signaling from ascaroside-sensing neurons, and DAF-16, which mediates DAF-2/IGF signaling. We hypothesize that DAF-12 cooperates with DAF-3 or DAF-16 in a DA-independent manner to activate and modulate the alternative HBL-1 downregulation program (Figure 5). Although the immediate action of DAF-12, together with DAF-3 or DAF-16, would likely be transcriptional, ultimately the alternative pathway for HBL-1 down regulation appears to be post-transcriptional, as we found that suppression of *daf-12(rh61)* or *mir-84(lf)* by L2d involves contributions from *lin-4* and *let-7 family* microRNAs, in addition to the miRISC cofactor NHL-2 ^28^ and also LIN-46, which is not thought to function as a transcriptional regulator ^29^.

Prior to this study, it was not clear whether the inhibition of dauer larva formation by exogenously-supplied DA reflects prevention of L2d in addition to a block of dauer-commitment. We observed that exogenous DA hormone at levels sufficient to prevent ascaroside-induced dauer formation does not prevent the extension of second larval stage development characteristic of L2d. Also, we observed that *daf-12(rh61)* animals, which are DA-insensitive and dauer-commitment defective nevertheless can readily undergo L2d. These observations suggest that L2d is initiated independently of DAF-12-DA activity, and are consistent with our model that a DA-independent function of DAF-12, together with activated DAF-3 or DAF-16, promote expression of the L2d program.

The studies described here involve animals that traverse the bi-potential L2d stage, but elect the option of resuming L3 development instead of dauer larva arrest. Previous studies showed that for animals that enter dauer diapause arrest and later are induced to emerge from dauer arrest and complete post-dauer L3 and L4 development, a similar change in the HBL-1 down-regulation program is evident^13^. Here we show that the genetic circuitry controlling cell fate progression via HBL-1 in larvae undergoing L2d development is similar to the circuitry previously identified for animals that arrest as dauer larvae ^13^. In particular, for both the L2d-to-L3 trajectory, and the L2d-to-dauer-to-postdauer trajectory, an enhance role for *lin-46* is evident, compared to the rapid continuous development trajectory. Moreover, *lin-4* and NHL-2 have similarly prominent roles in both the L2d-only and the post-dauer trajectories. These findings, that the rewiring of HBL-1 down regulation for the L2d-only trajectory is largely similar to that for dauer-postdauer trajectory, is consistent with the fact that dauer larvae arrest is always preceded by L2d. Therefore, the rewiring associated with traversing post-dauer likely is initiated during L2d. The observation that suppression of the retarded phenotypes of certain mutants is more potent for the L2d-dauer-postdauer trajectory compared to the L2d-only trajectory (Figure 2), suggests that rewiring may be partially implemented during L2d, and more fully engaged in association with dauer commitment (Figure 5).

Interestingly, there is evidence that the L2d component of the alternative HBL-1 down regulation program may be dosage sensitive. The length of the L2d stage is reported to be variable, depending on the strength of the dauer-inducing conditions ^14^. Similarly, we observed that larval development of *daf-12(rh61)* was delayed the most by ascr#5 compared to ascr#2 or ascr#3 (data not shown), and ascr#5 also elicited the strongest suppression of *daf-12(rh61)* phenotypes (Supplemental Figure 3A). This suggests a correlation between the length of the L2d and the degree of the suppression of *daf-12(rh61).* It appears that selection of the L2d trajectory, which occurs before the L1/L2 molt (Figure 1), may not be in itself sufficient for full suppression. These observations suggest that the degree to which L2d rewiring is implemented may depend on an integration of stress signals experienced by the larva during the L2d. Consistent with a quantitative gradient of rewiring, perhaps linked to the degree of stress experienced by each animal, the distributions of phenotypes in a population of suppressed *daf-12(rh61)* animals can range broadly from partial suppression to near complete suppression (For example: Supplemental figure S3A, rows 8 and 9).

Our observations of the kinetics of HBL-1 down regulation during the L2 and L2d stages suggest that HBL-1 levels are maintained at relatively high levels for much of the L2 or L2d stage, and down-regulated near the end of the stage (Figure 6 and Supplemental figure S4). This suggests that down-regulation of HBL-1, and hence commitment to L3 cell fate specification, may be gated by some event(s) coupled to the completion of the L2 or L2d stage. This would be consistent with a model wherein most of the L2d is occupied with rewiring of pathways upstream of HBL-1, followed by implementation of HBL-1 down regulation at the end of the L2d, in association with commitment to the L3 cell fate.

In summary, we show that environmental and endocrine stress signals that regulate developmental progression and alternative developmental trajectories rewire the genetic regulatory network controlling temporal cell fate transitions during larval development of *C. elegans.* Our findings provide insight into how the regulation of temporal cell fate transitions can be adapted to accommodate stressful conditions that challenge developmental progression. Coordinate regulation of developmental progression and temporal cell fates seem to confer elasticity to *C. elegans* development, wherein the proper developmental schedule of cell fate transitions is unaffected by variations in developmental rate or developmental trajectory.

*C. elegans* larvae possess other life history options, in addition to the delayed developmental progression associated with the L2d-dauer trajectory. For example, appropriate pheromone signals and nutritional status can result in an acceleration of *C. elegans* development ^31-33^. Moreover, larvae can temporarily suspend developmental progression at specific checkpoints in the late L2, L3, or L4 stages in response to acute starvation ^34^. We hypothesize that these signals that either accelerate development or result in programmed developmental arrest would also be associated with appropriate rewirings of the genetic regulatory pathways regulating temporal cell fate progression; so that cell fate transitions are appropriately synchronized with the dynamics of larval stage progression. Lastly, rewiring mechanisms induced by environmental signals, such as that described here, might also be adapted for the evolution of morphological plasticity or polyphenism observed in certain nematodes ^35,36^ and other animals ^37^.

## METHODS

### *C. elegans* culture conditions

*C. elegans* strains used in this study are listed in Table S1. *C. elegans* was cultured on nematode growth media (NGM) using standard techniques ^38^ and fed with the *E. coli* HB101 strain.

For experiments involving the administration of ascarosides and/or DA, we adopted the protocol described by Butcher *et. al.* ^17^ with modifications. *C. elegans* was fed with the *E. coli* OP50 strain on plates containing 3 mL of 1% agarose (SeaKem® LE agarose, Cat#50004) with nematode growth media (NGM) without peptone. Synthetic ascarosides (kindly provided by the labs of Frank Schroeder and Jagan Srinivasan) were dissolved in ethanol (stock concentrations: ascr#2 5.69 mM, ascr#3 3.81 mM, and ascr#5 4.09 mM) and added to the melted agarose prior to plate-pouring to achieve the desired final concentration (3 µM, if not specified). Plates were seeded the next day with *E. coli* strain OP50 as follows: OP50 was grown in liquid Luria Broth (LB) media until the culture reached OD600=0.6-0.7. Then, the bacterial culture was pelleted by spinning at 3500 rpm for 10 minutes. The pellet was washed twice with a volume of sterile water equal to the LB culture volume. Finally, the pellet was resuspended in a volume of sterile water equal to one-fifth of the initial LB culture volume. 50 µLs of this washed and 5x concentrated OP50 culture were used to seed ascaroside plates. To prepare ascaroside plates also containing Δ4-dafachronic acid (DA; prepared from Cayman Chemical, item no. 14100, by dissolving 1 mg of DA in ethanol) 50 µL of water containing DA at specified concentrations was added onto the lawn of bacteria.

### Analysis of extra seam cell phenotypes

Gravid adult animals were bleached to isolate eggs. Eggs were placed on control or treatment plates and cultured at 20°C unless otherwise specified. The worms were scored at the young adult stage for the number of seam cells using fluorescence microscopy with the help of the *maIs105 [pCol-19::gfp]* transgene that marks the lateral hypodermal cell nuclei or the *wIs51[pScm::gfp]* transgene that marks the seam cell nuclei.

### Sample collection and Taqman assay for microRNA quantification

Synchronized L1 larvae of *daf-12(rh61)* were raised on control or ascr#2-3-5 plates at 20°C. Larvae that reached the L2/L2d-to-L3 molt were identified and picked under a dissecting microscope and collected in M9 media within two hours. For each experimental condition, three biological samples were collected, containing approximately 200 larvae per sample. Collected worms were snap-frozen and kept at −80°C until RNA extraction. RNA was extracted using the Trizol reagent (Invitrogen).

Two microliters of 30 ng/μL RNA samples were used for reverse transcription, and multiplex miR-Taqman reactions were carried out according to the manufacturer’s instructions, and using an ABI 7900-HT Fast-Real Time PCR System (Applied Biosystems). MicroRNAs were assayed in three technical replicates for each biological sample. Five highly expressed microRNAs that have not been reported to be environmentally regulated (*mir-1, lin-4, mir-52, mir-53, and mir-58*), were used as a control microRNA set. The average Ct of the control microRNA set was used to normalize the Ct’s obtained from *let-7* family microRNAs.

### Tagging of *hbl-1* at its endogenous locus using CRISPR/Cas9 genome editing

A mixture of plasmids encoding Cas9 (pOI90), and single guide RNAs (sgRNAs) targeting the site of interest (pOI89) and the *unc-22* gene (pOI91) as co-CRISPR marker^39^, a donor plasmid (pOI191) containing the mScarlet-I sequence^40^ flanked by homology arms, and a *rol-6(su1006)* containing plasmid (pOI124) as co-injection marker was injected into young adult germlines. F1 roller and/or twitcher animals were cloned and screened by PCR amplification for the presence of the expected homologous recombination (HR) product. F2 progeny of F1 clones positive for the HR-specific PCR amplification product were screened for homozygous HR edits by PCR amplification of the locus using primers that flanked the HR arms used in the donor plasmid. Finally, the genomic locus spanning the HR arms and mScarlet DNA was sequenced using Sanger sequencing. A single worm with a precise HR edited locus was cloned. The allele is named as *ma430* [*hbl-1::mScarlet-I*] and it is backcrossed twice. The ma430 allele contains the following the DNA sequence (underlined) inserted in-frame with the *hbl-1* coding sequence before the stop codon (bold): “…CCACATGTACCAAGCCAGACACCAA<u>tctggaggtggatctggaggtggatctggaggtggatctGTCAGCAAGGGAGAGGCAGTTAT</u> <u>CA…<-- rest of the mScarlet-I -->…CGGAGGAATGGACGAGCTCTACAAG</u>**TAA**TGA*GG*ACGTCCTCGTTAAGGAA…”

#### Cloning of sgRNA plasmids

All plasmids in the injection mix had the same plasmid backbone which was derived from pRB1017^41^. sgRNA encoding plasmids were derived from pRB1017 (first to generate pOI83) by modifying the tracr encoding sequence to (F+E) form of the tracr ^42^, which was reported to increase the CRISPR efficiency in *C. elegans* ^43^. pOI89 and pOI91 sgRNA encoding plasmids were sub-cloned from pOI83 by ligating annealed primer pairs to the BsaI cloning sites ^41^. The following primers were annealed and cloned into pOI83 to generate pOI89 and pOI91:

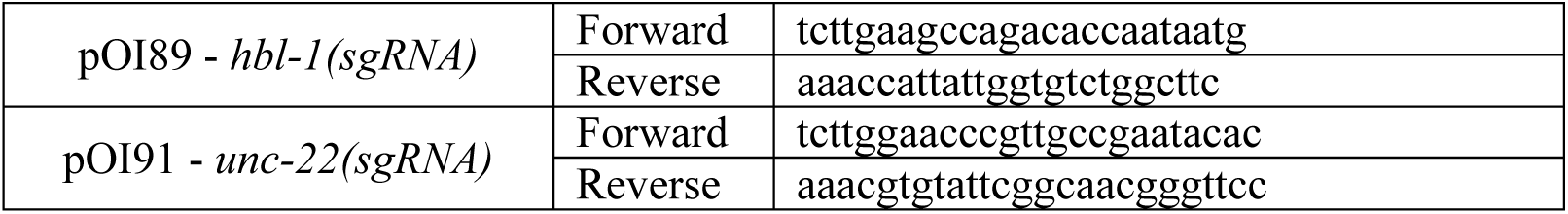

#### Cloning of pOI191 HR Template plasmid

The Golden Gate Assembly Kit (NEB Cat#E1600) is used to fuse two PCR fragments: *mScarlet* (PCR amplified from pSEM91^44^) and a DNA fragment containing left and right HR arms fused to the pRB1017 plasmid backbone (PCR amplified from pOI115, which was previously generated by assembling four PCR fragments containing a left HR arm, a GFP, a right HR arm, and the backbone of pRB1017). Single colony purified plasmid DNA was used to identify colonies containing the precise assembly of the HR arms and mScarlet. pOI191 was derived from this mScarlet clone (pOI186) by mutating a single nucleotide to convert mScarlet to mScarlet-I using the Q5 Site-Directed Mutagenesis kit (NEB Cat#E0554). Primers used for cloning:

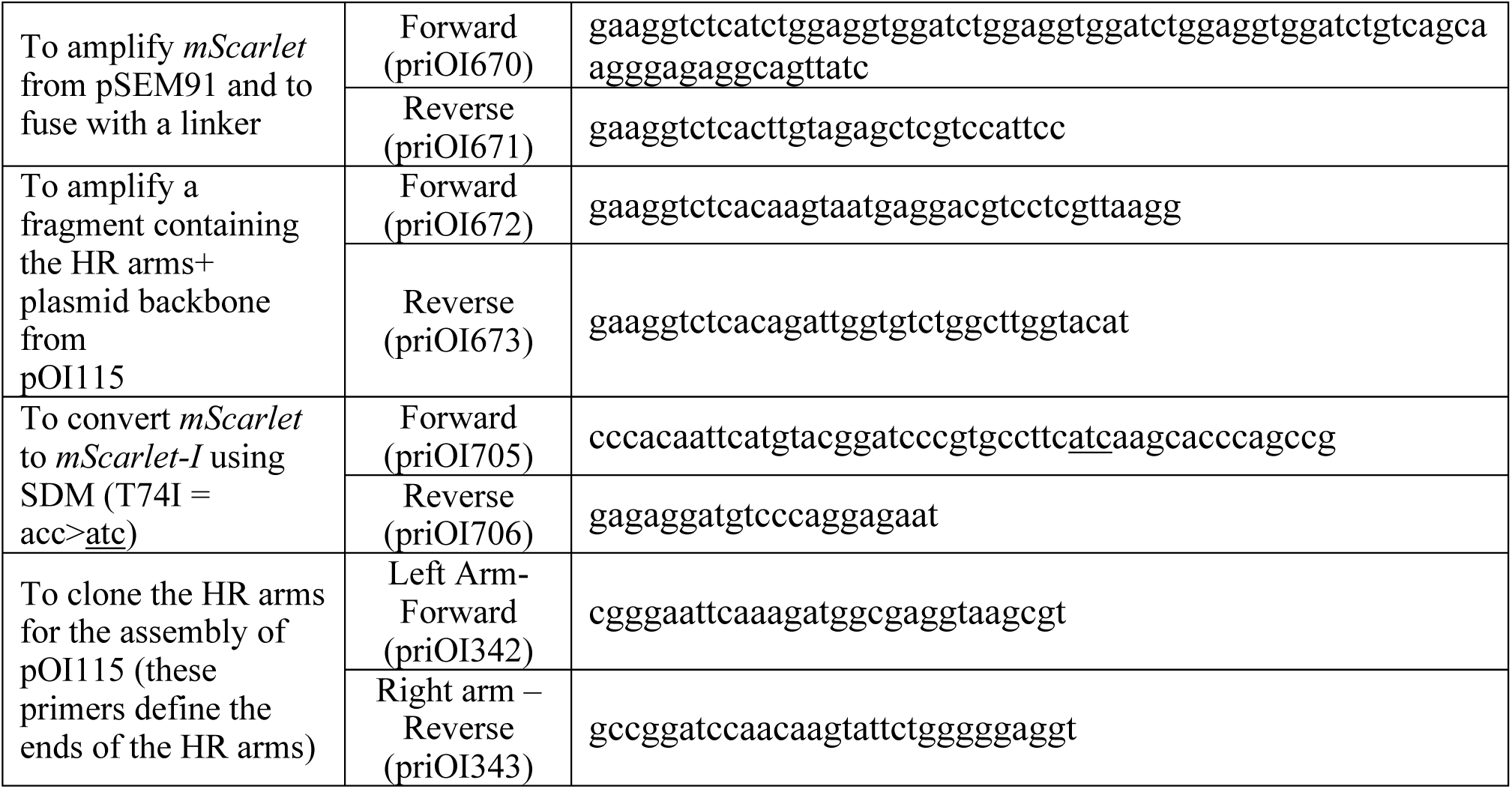

## SUPPLEMENTAL INFORMATION

**Supplemental Figure S1. Related to Figure 2.**
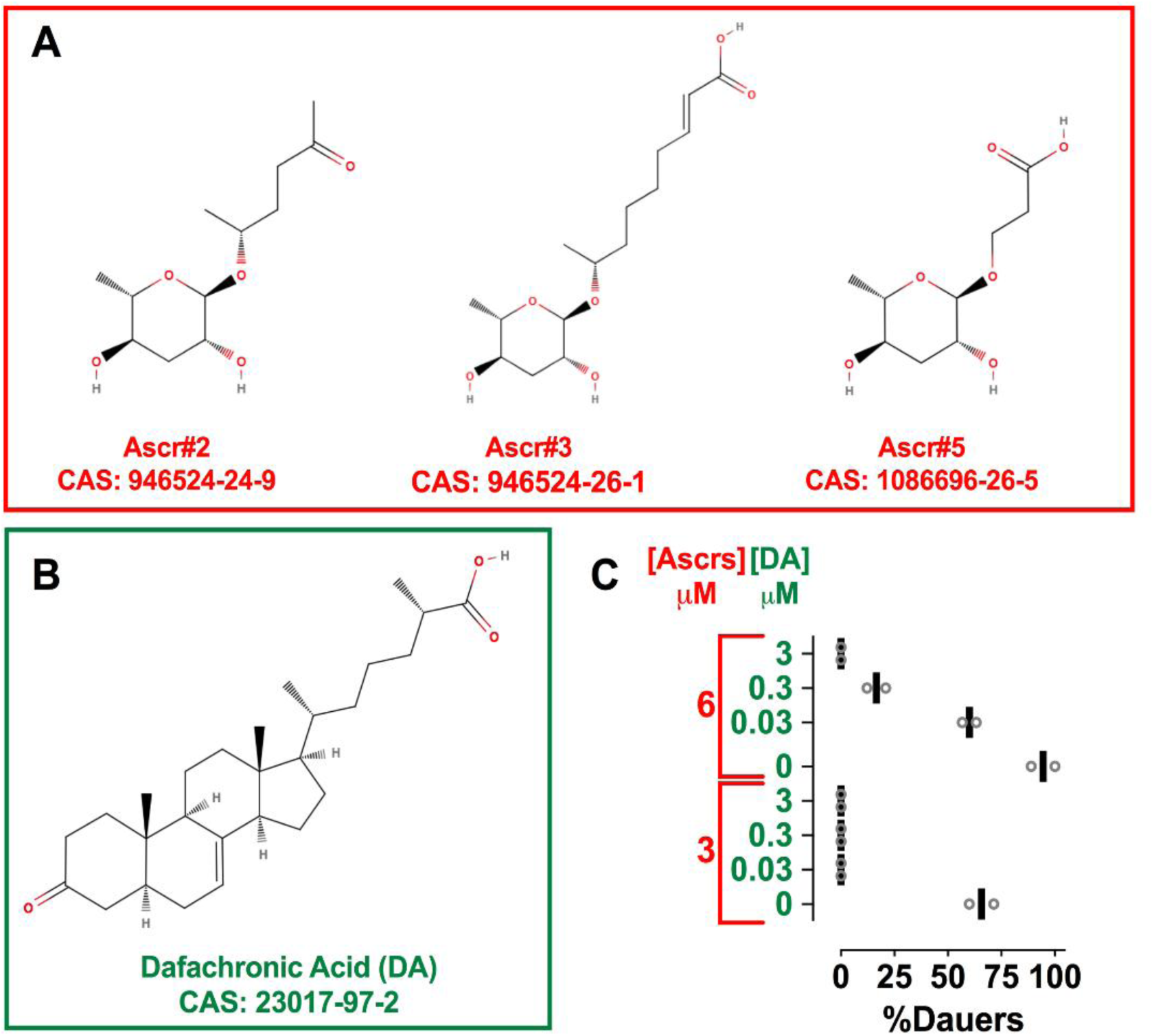
Chemicals and conditions used to obtain worm populations developing continuously through L2d trajectory. **A-B)** Molecular structures (using www.molview.org), names, and CAS numbers of the chemicals used in the ascaroside (dauer inducing), and ascaroside plus dafachronic acid (both dauer inducing and dauer commitment inhibiting) plates. **C)** Two different concentrations of Ascrs assayed in combination with four different concentrations of DA to determine conditions that prevent dauer formation in the presence of ascarosides. Percent dauer formation in the presence of different combinations of ascarosides (Ascrs: equimolar mixture of ascr#2, ascr#3, and ascr#5) and dafachronic acid (DA) are plotted. DA inhibits dauer commitment but not L2d, which is evident by slowing of larval development. The combination of 3 µM of Ascrs and 0.03 µM DA was used as the L2d-inducing (Ascrs+DA) condition to test the effect of the L2d trajectory on the retarded heterochronic phenotypes of *mir-48/84/241* in Figure 2.

**Supplemental Fig S2. Related to Figure 2.**
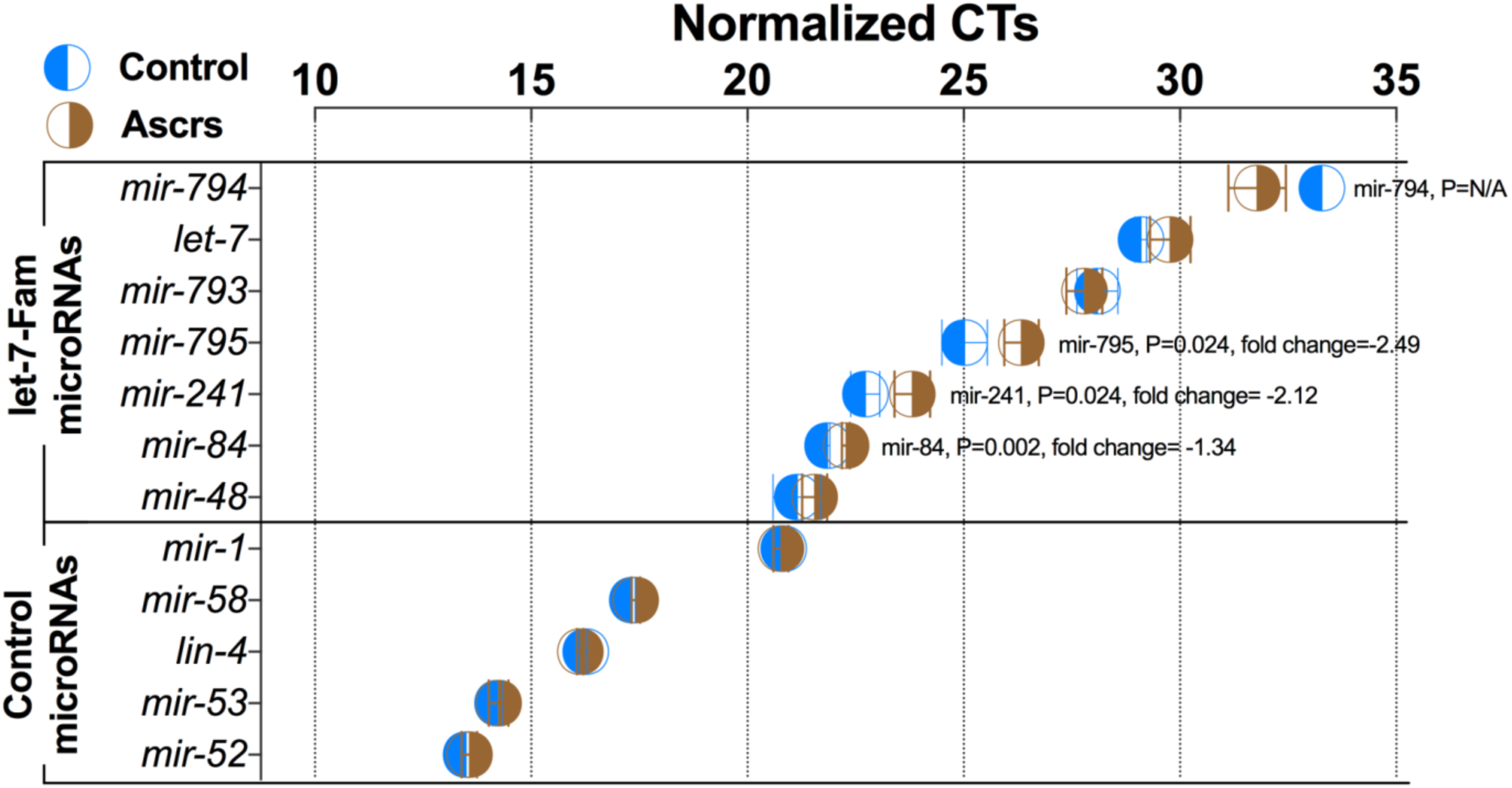
Ascarosides do not result in an increase in *let-7* family levels in *daf-12(rh61)* background. *let-7* family microRNAs in L2-to-L3 (control) vs. L2d-to-L3 (Ascrs) molting larvae of the *daf-12(rh61)* mutant were quantified using Taqman assays as described in the material and methods section. MicroRNAs that are highly expressed and not environmentally regulated were used as the normalization set (Control MicroRNAs). The expression levels of three *let-7* family microRNAs (*mir-84, mir-241, mir-795*) were slightly but statistically significantly reduced in the presence ascarosides (L2d-to-L3 molt). This reduction of *let-7* family levels is in contrast with the observed suppression of retarded heterochronic phenotypes of *daf-12(rh61)* in the presence of ascarosides. The lack of an upregulation of *let-7* family microRNAs in the presence of ascarosides is in line with the idea that an alternative, *let-7*-independent, mechanism is responsible for the ascaroside-mediated suppression of the heterochronic phenotypes. *mir-794* was not detected in two biological control samples (presumably due to low expression level); therefore, we do not know if there is a statistically significant up-regulation of *mir-794* in the presence of ascarosides.

**Supplemental Fig S3. Related to Figure 3.**
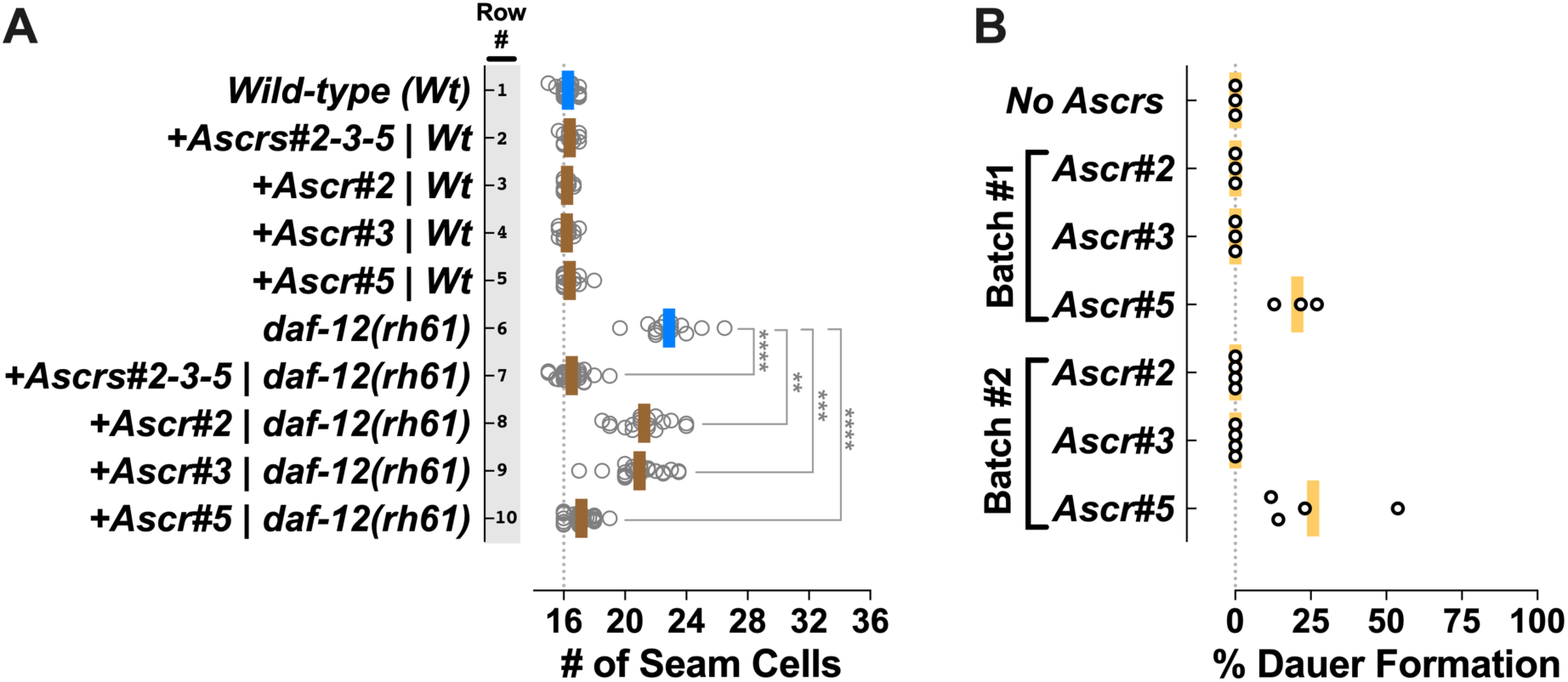
**A)** Individual components of the pheromone cocktail can suppress the extra seam cell phenotypes of *daf-12(rh61)* and ascr#5 is the most potent suppressor. Number of seam cells in young adult animals cultured under different ascaroside conditions. Each dot in the plot shows the number of seam cells of a single young adult animal, and solid lines (blue: rapid trajectory; brown: delayed [l2d] trajectory) indicate the average seam cell number of the animals scored for each condition. Ascrs#2-3-5 plates contained all three ascarosides at 3 µM final concentration of each ascaroside mixed in NGM-agarose media. Ascrs#2, Ascr#3, and Ascr#5 plates contained 3 µM final concentration of each ascaroside mixed in NGM-agarose media. The student’s t-test was used to calculate statistical significance (p): n.s.(not significant) p>0.05, *p<0.05, **p<0.01, ***p<0.001, ****p<0.0001 **B) Ascr#5 alone can induce dauer formation**. At 3.3 µM concentration, using agarose NGM plates with no peptone, seeded with washed and concentrated *E. coli* OP50 culture as described in the materials and methods, Ascr#5 alone was sufficient to induce dauer formation but not Ascr#2 or Ascr#3. We tested single ascarosides in two different batches of plates and using three or four replicates. Both experiments were performed at 20°C. Each dot on the plots shows percent dauer formation on a single plate. We maintained population sizes small (<55 worms per plate) and comparable across different Ascr plates to minimize the potential effect of the accumulation of ascarosides secreted by the worms on the plates.

**Supplemental Figure S4. Related to Figure 7.**
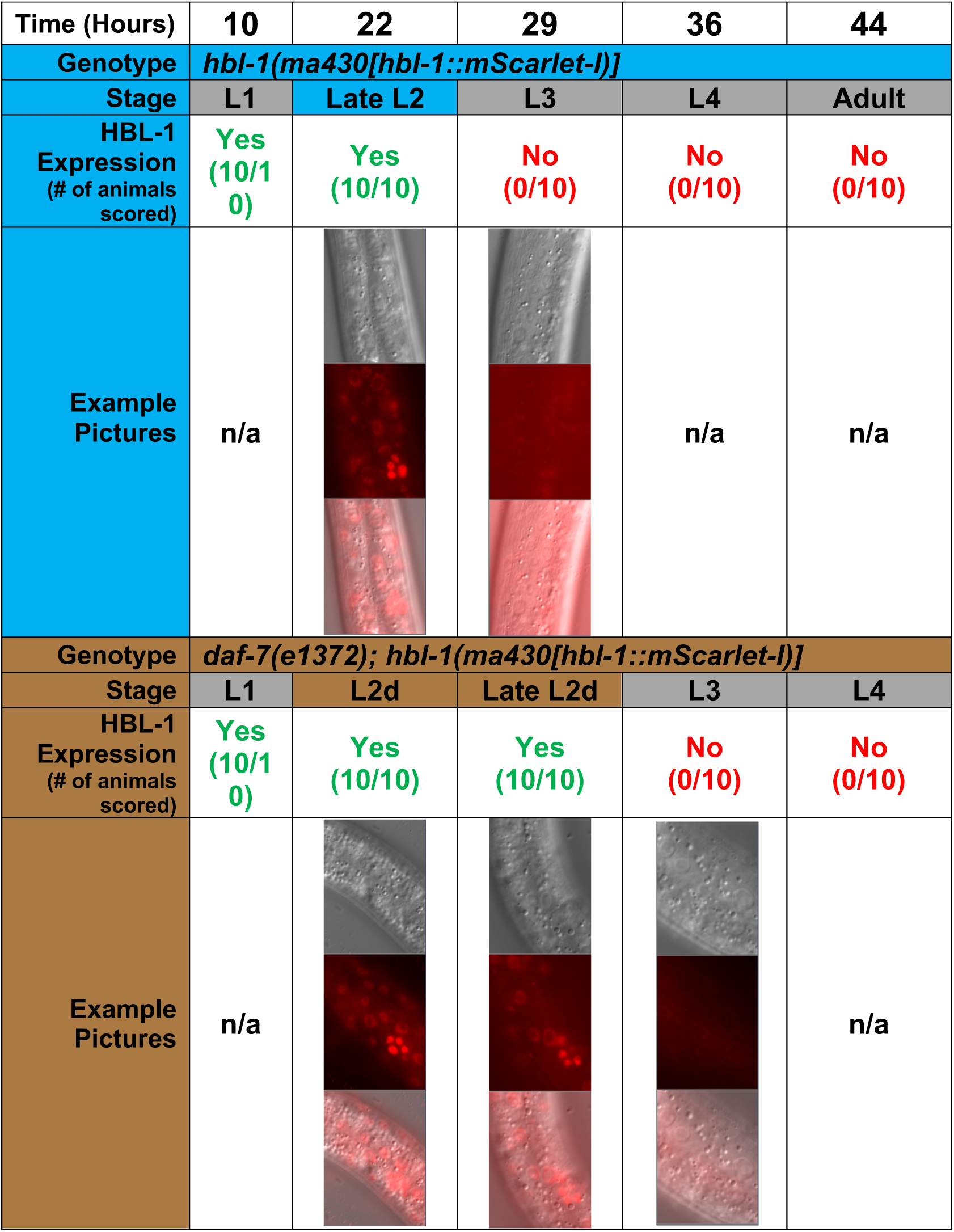
HBL-1 is present throughout the L2 or the lengthened L2d stage but it is absent at the post-L2 or post-L2d L3 stage. Endogenously tagged HBL-1 expression was examined in wild-type and *daf-7(e1372)* backgrounds at 20°C. *daf-7(e1372)* animals form dauer larvae at 25°C but they develop continuously going through L2d at 20°C. For each time point and genotype ten animals were examined. In all animals at 22 hours, also in *daf-7(e1372)* animals at 29 hours, L2 stage specific V5p cell divisions were observed to have occurred, indicating that the animals were at a late phase of the L2 stage (in L2d animals the lengthening of the second larval stage appeared to occur after the execution of these L2 stage cell divisions). HBL-1 is detected in late L2 and L2d larvae but it was not detected in post-L2 and post-L2d L3 stage larvae, indicating that the duration of HBL-1 expression is lengthened as the L2 stage is lengthened during L2d, and in both post-L2 L3 and post-L2d L3 animals HBL-1 is downregulated.

**Supplemental Table S1.**
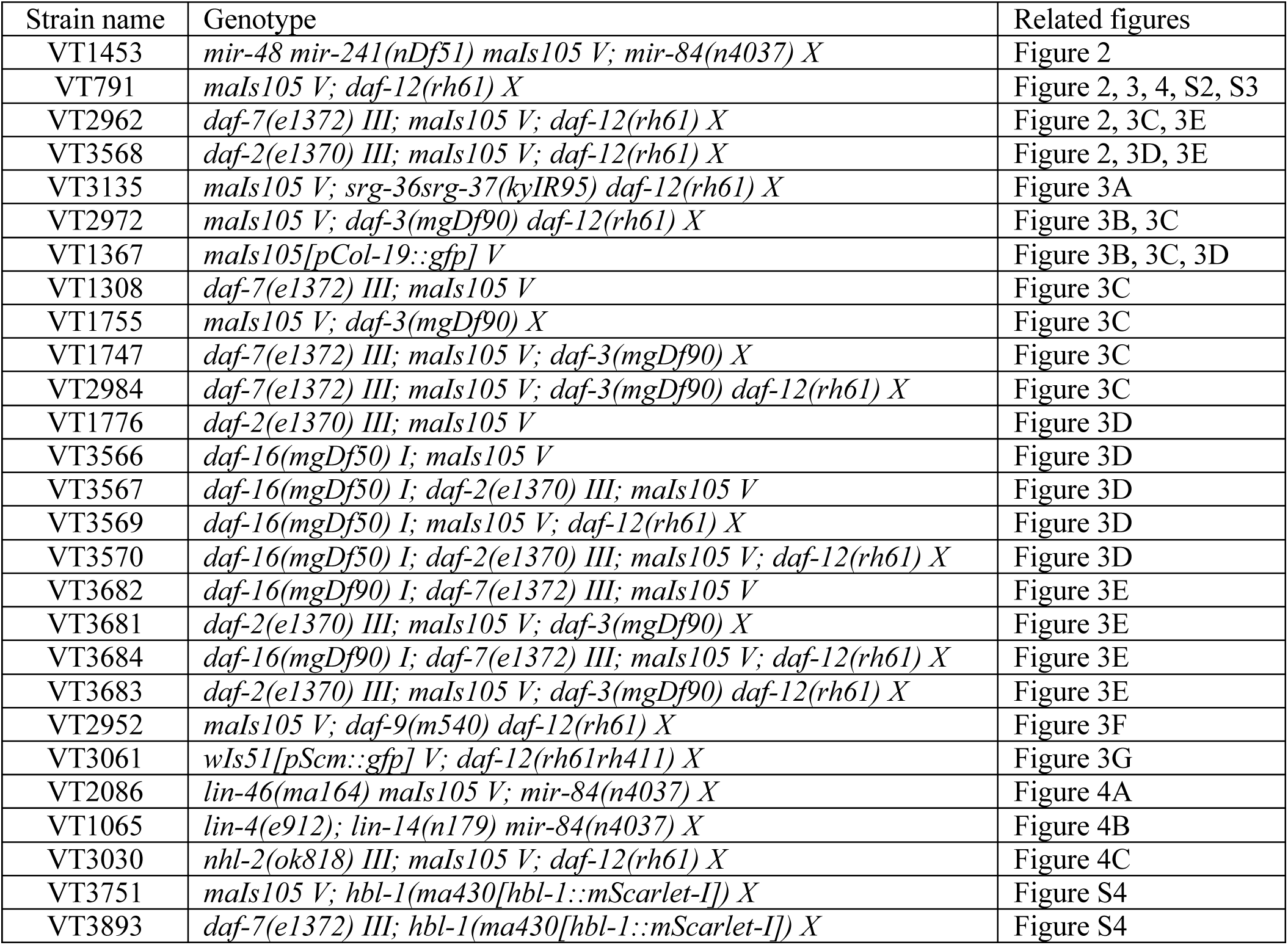
*C. elegans* strains used in this study.

